# Sexual reproduction as bet-hedging

**DOI:** 10.1101/103390

**Affiliations:** Department of Evolutionary Biology and Environmental Studies, University of Zurich, Winterthurerstrasse 190, 8057 Zurich, Switzerland; Evolution and Ecology Research Centre, School of Biological, Earth and Environmental Sciences, University of New South Wales, Sydney, New South Wales 2052, Australia

**Keywords:** Bet-hedging, Environmental fluctuation, Evolutionary games, Geometric mean fitness, Sexual reproduction

## Abstract

In evolutionary biology, bet-hedging refers to a strategy that reduces the variance of reproductive success at the cost of reduced mean reproductive success. In unpredictably fluctuating environments, bet-hedgers benefit from higher geometric mean fitness despite having lower arithmetic mean fitness than their specialist competitors. We examine the extent to which sexual reproduction can be considered a type of bet-hedging, by clarifying past arguments, examining parallels and differences to evolutionary games, and by presenting a simple model examining geometric and arithmetic mean payoffs of sexual and asexual reproduction. Sex typically has lower arithmetic mean fitness than asex, while the geometric mean fitness can be higher if sexually produced offspring are not identical. However, asexual individuals that are heterozygotes can gain conservative bet-hedging benefits of similar magnitude while avoiding the costs of sex. This highlights that bet-hedging always has to be specified relative to the payoff structure of relevant competitors. It also makes it unlikely that sex, at least when associated with significant male production, evolves solely based on bet-hedging in the context of frequently and repeatedly occupied environmental states. Future work could usefully consider bet-hedging in open-ended evolutionary scenarios with *de novo* mutations.

## Introduction

Evolutionary dynamics in natural populations are under the combined effect of directional selection and randomness that comes from various sources, including environmental fluctuations and demographic stochasticity. Accurate predictions of evolutionary dynamics depend, in principle, on all the moments of the fitness distribution of individuals and their relative weights. In general, populations tend to be driven towards phenotypes that maximise the odd moments (mean fitness being the first moment) while minimising the even moments of their fitness distributions (variance being the second moment) (Rice, 2008). This implies that the adverse change of one moment can potentially be compensated by the beneficial changes of other moments. Most attention has been placed on the possibility that decreased mean fitness might be sufficiently compensated for by a concominant decrease of the variance in fitness, such that the strategy with diminished mean fitness out-competes others over time (Philippi and Seger, 1989). Because strategies that gain success by manipulating fitness variance intuitively fit the idea of “hedging one’s bets” (Starrfelt and Kokko, 2012), this has given rise to a precise biological meaning of the phrase “bet-hedging” (Slatkin, 1974): it refers to strategies that have diminished arithmetic mean fitness, but also reduced variance (and are often studied with the aid of geometric mean fitness).

Bet-hedging bears some similarity to mixed strategies in evolutionary games (the phrase “optimal mixed strategies” (Haccou and Iwasa, 1995, 1998) has been used near-synonymously with bet-hedging under non-game-theoretical contexts): some forms of bet-hedging imply the production of different kinds of offspring (e.g. different sizes of tubers in the acquatic macrophyte *Scirpus maritimus*, (Charpentier et al., 2012)). Although both bet-hedging and mixed strategies (in game theory) can lead to a mix of phenotypes in the population, there are two important differences between the concepts: first, the adaptive reasoning is different, and second, bet-hedging can also occur without phenotypic variation. To explain the first difference: In evolutionary games, the payoff of an individual depends on the action of other individuals in the population. This is not a requirement in bet-hedging, where the payoff is typically thought to be determined by the stochastically varying environment (though, as our examples show, others’ presence can matter too: e.g. sexual reproduction to diversify one’s offspring to cope with environmental change would not work if diversity has been lost). A typical context in which bet-hedging is discussed is rainfall that varies over time (Seger and Brockmann, 1987; Starrfelt and Kokko, 2012). Under such conditions it can then be beneficial if an individual can produce both wet-adapted and dry-adapted offspring, so that regardless of the conditions in a given year, some fraction of offspring will survive; a non-bet-hedger’s entire genetic lineage might disappear as soon as an environmental condition occurs to which it is not adapted.

The second difference between mixed strategies and bet-hedging is that the latter can work without there being a “mix” of any kind. Instead of diversifying offspring, a so-called *conservative* way of bet-hedging is to produce only one type of offspring that performs relatively well under all different environments, while not being the best under any of them (“a jack of all trades is the master of none”). This can also reduce fitness variance, and qualify as bet-hedging if it is achieved at the cost of reduced mean fitness.

One prominent example that seems to have the characteristics of bet-hedging, but is less often mentioned in a bet-hedging context, is sexual reproduction, where offspring are formed using genetic material from two parents (because nature is diverse there are definitional complications and grey zones regarding what counts as sex; see Lehtonen and Kokko (2014)). Producing offspring in this way, as opposed to the simpler option of asexual reproduction, incurs costs in many different ways (reviews: Lehtonen et al. (2012); Meirmans et al. (2012)). The best known cost, and the one we focus on here, is the two-fold cost of males: if the offspring sex ratio is 1:1 and males and females are equally costly to produce, a mother will use 50% of her resources on offspring that do not themselves contribute material resources to the next generation (Maynard Smith, 1978), and this slows the growth of sexual populations compared with asexual ones. Consequently, sexual reproduction – when it involves producing males – is expected to lead to a reduction of mean fitness. But on the other hand, through mixing genetic material from different lineages, sex provides a potent way of producing offspring whose genomes differ from each other. If some always do well no matter what the state of the environment, the variance of reproductive fitness can be reduced compared with an asexual lineage.

Given that effects on genetic diversity are central and much discussed in the sex literature (e.g. Hartfield and Keightley (2012)), it is surprising that the biological literatures on bet-hedging and on sex are relatively separate. Mixed strategies have been shown to be advantageous in a fluctuating environment (Haccou and Iwasa, 1995, 1998; McNamara et al., 1995). Haccou and Iwasa (1995) have shown that the optimal strategy can involve bet-hedging under a fluctuating environment in unstructured populations, and showed how to calculate the strategy explicitly for a given payoff function and a given distribution of the environmental parameters. In addition, the optimal bet-hedging strategy is robust against small perturbations of the distribution of environmental conditions and/or the payoff function (Haccou and Iwasa, 1998). Cooperative games between kin can also help maximise the geometric mean fitness of species in fluctuating environments (McNamara, 1995). Furthermore, the strategy that maximises the geometric mean fitness is more likely to evolve in species of non-overlapping generations compared to species with substantial parental survival. In the latter case, the strategy that maximises the arithmetic mean fitness is more likely to evolve (Haccou and McNamara, 1998). The review of Grafen (1999) discusses different ways of optimising reproductive fitness in a fluctuating environment. None of these studies, however, have explicitly pointed out that sexual reproduction can be a form of bet-hedging.

Williams (1975) in his classic book on sex discusses a “lottery model” using the verbal analogy of buying ever more copies of the same number on a lottery ticket (asexual reproduction) vs. buying fewer but a more diverse set of numbers (sexual reproduction). The analogy to a real-life lottery is not perfect, in the sense that asexually produced offspring are often not totally redundant copies of each other, i.e. they do not necessarily have to share the prize if both have a winning number: two asexually produced offspring usually leave more descendants than just one, especially if they disperse to different localities and no longer compete for the same resources ((Williams, 1975) p.16). The correspondence between Williams’ lottery model and bet-hedging, on the other hand, appears perfect. But Williams (1975) did not use explicit bet-hedging terminology, probably because it had only very recently been imported to evolutionary terminology (Slatkin, 1974).

Williams (1975) emphasised the need to consider the spatial arrangement of offspring to determine whether, e.g., 10 “winning tickets” can win 10 prizes, which requires dispersal to avoid competition with relatives, or are expected to win less ((Williams, 1975) p.53). The emphasis in Williams’ idea is that the winning numbers vary over time (but not necessarily over space). In a context where dispersal is limited, a similar idea has been formulated emphasising resource diversity rather than its temporal fluctuations. The relevant metaphor is a “tangled bank”, a rather poetic phrase that has its origin in Darwin’s *On the origin of species*. Darwin contemplated “a tangled bank, clothed with many plants of many kinds, with birds singing on the bushes, with various insects flitting about, and with worms crawling through the damp earth…” (Darwin, 1859). Darwin was not talking specifically about sex, but about life and its evolution in general. Nevertheless, the “tangled bank” has since acquired a specific meaning (Bell, 1982), becoming a metaphor of genetic polymorphisms favoured in environments that might not vary much temporally but that, based on diverse resources present at the same site, offer multiple niches and the resultant higher total carrying capacity for different phenotypes as a whole (“the environment is now more fully utilised…, the carrying capacity of the diverse population will inevitably exceed that of either single clone.” (Bell, 1982) p.130). In the “tangled bank” scenario, the carrying capacity of each single clone depends on the distribution of different niches in the environment. The carrying capacity of the entire diversified population in the heterogeneous environment is larger than any of the single clones.

Although the “tangled bank” does not require a temporally fluctuating environment, the diversity of different clones is maintained better if the environment changes frequently (Bell, 1982). In addition, in the “tangled bank”, the fitness of a single clone depends not only on the abundance of different niches, but also is frequency-dependent when competing for the same niche or invading a new niche (Bell, 1982). Therefore, the “tangled bank” may capture aspects of the benefits of sexual reproduction, but it does not perfectly correspond to bet-hedging.

## 2 Bet-hedging via heterozygotes and sexual reproduction

We examine in the following the conditions under which sexual reproduction might spread as a form of bet-hedging. Our model considers a large well-mixed population where a proportion s of the young produced are male. Note that our assumption of large (infinite) population size allows us to focus on the effects of environmental stochasticity without confounding effects of demographic strochasticity. Asexual individuals are all female. The adaptation to the amount of rainfall in the environment is determined by a diploid genetic locus that has two alleles. The AA genotype is well adapted to the wet environment, whereas the aa genotype is dry-adapted. The heterozygote Aa has intermediate fitness in both environments, but not necessarily exactly the mean of aa and AA. Example fitness values for each genotype under different environments are show in matrix (1).

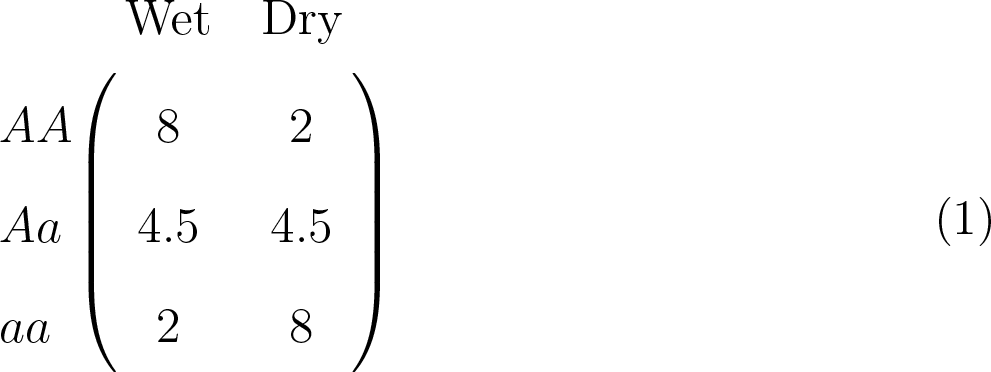

Consider a case where wet and dry environments occur at equal frequencies, and all individuals are asexual females. Table 1 shows the arithmetic mean and geometric mean fitness of the different asexual types. The heterozygote (Aa) has the lowest arithmetic mean fitness, but the highest geometric mean fitness, which predicts higher evolutionary success if we ignore higher moments of the fitness distribution (Starrfelt and Kokko, 2012). The asexual heterozygotic form becomes thus a bet-hedging strategy when compared with the two other asexual homozygotic forms. This form of bet-hedging is *conservative*: all Aa individuals have the same expected fitness under both environmental conditions.

**Table 1:** The payoff structure under wet and dry years: the arithmetic mean (AMean) and the geometric mean (GMean) of the payoffs of asexual lineages, as well as of a sexual population assumed to be at the Hardy-Weinberg equilibrium.

|  | Wet | Dry | AMean | GMean |
| --- | --- | --- | --- | --- |
| asex-AA | 8 | 2 | 5 | 4 |
| asex-Aa | 4.5 | 4.5 | 4.5 | 4.5 |
| asex-aa | 2 | 8 | 5 | 4 |
| sex-population | $4.75(1-s)$ | $4.75(1-s)$ | $4.75(1-s)$ | $4.75(1-s)$ |

In contrast to the conservative approach of the asexual heterozygotes, the sexual population as a whole can also be seen to bet-hedge, in this case by producing offspring of different genotypes. It is therefore of interest to ask if sex is a bet-hedger with respect to AA, Aa, aa or perhaps all of them. The comparison is more complicated than the above one, not only because sex produces young that differ from each other (and thus differ in the long-term growth rate impacting the original parent’s contribution to the future gene pool), but also because the frequencies of genotypes in the offspring of any given parent depend on the genetic composition of the population as a whole – which in turn depends on how selection has worked on it in the recent past: a run of wet years will have favoured the A allele, dry years do the opposite.

We initially assume that the sexual population is always under Hardy-Weinberg equilibrium (Hardy, 1908; Weinberg, 1908) and that the two alleles are equally abundant. This is a strong assumption that is expected to be violated as soon as selection is applied, but we nevertheless consider it as a useful thought experiment, because the genetic background that an allele faces is then constant across generations (genotypic proportions are always expected to be *x*_AA_ = 1/4, *x*_Aa_ = 1/2, and *x*_aa_ = 1/4). Given that only females contribute directly to offspring production (males only impact the genetic diversity of young she produces), the expected growth rate of the sexual population equals (8/4 + 4.5/2 + 2/4)(1 − *s*) = 4.75 (1 − *s*), where *s* is the proportion of males. If the sexual population achieves this growth rate in every year (which requires that it maintains itself at the Hardy-Weinberg equilibrium), and as long as *s* is not too large, it has performed perfect bet-hedging as the geometric mean now equals the arithmetic mean, which is its maximum value.

But is this geometric mean fitness higher than that of the specialist asexuals (AA and aa)? The answer depends on the cost of sex, which we here model as the proportion s of offspring developing as males. Sex beats AA or aa asexual genotypes if *s* < 0.158, while beating the bet-hedging asexual genotype (Aa) is harder: it only occurs if *s* < 0.0526.

While the example shows that sexual reproduction can, in principle, be a bet-hedging strategy, it simultaneously shows how difficult it is for sex to evolve based on this benefit alone, especially if competing against asexual types that also bet-hedge (conservatively). The cost of males is captured by *s*, and the more females produce sons, the higher this cost. Why males exist is a separate evolutionary conundrum from why sex exists: the alternative that is relevant for the “why males?” question is still sex, but without having some individuals specialise in the male strategy that fails to contribute directly to population growth. This question has its own set of game-theoretical answers (Bulmer and Parker, 2002; Lessells et al., 2009; Lehtonen and Kokko, 2011); the short summary is that (1) males can invade sexual populations despite the reduced growth rate, (2) their existence increases the vulnerability of sexual populations to invasion by asexuals, (3) if a population only consists of (sexual) females and males, sex ratios evolve to *s* = 0.5 under quite general conditions (West, 2009).

In Table 1, the arithmetic mean decreases rapidly with an increasing production of males, and any primary sex ratio greater than 15.8% males leads to sexuals being unable to resist invasion by any of the asexual options. Because male presence typically leads to much higher sex ratios, sex is unlikely to persist due to its bet-hedging benefits alone, at least in the simplistic setting of Table 1.

Sexual populations can resist invasions somewhat better (i.e. up to a larger fraction of sons produced) if the dimensionality of bet-hedging increases (i.e. it involves multiple traits). For example, besides the A/a locus that determines an individual’s fitness in response to the amount of rainfall, consider another diploid locus that impacts the adaptedness to high or low temperatures. Assume that an individual of the BB genotype is hot-adapted, an individual of the bb type is cold-adapted, and the Bb genotype is intermediate. Also assume the payoff matrices for rainfall and temperature adaptation has the same structure:

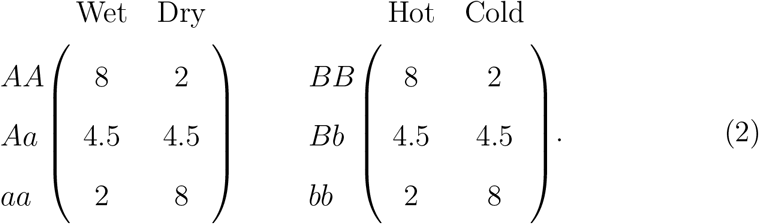

If different traits interact multiplicatively to determine the final fitness, then an AABB individual has payoff of 64 if the environment is both wet and hot (WH), 16 if the environment is wet but cold (WC), or dry but hot (DH), and 4 if the environment is both dry and cold (DC). Table 2 gives the complete list of payoffs of different genotypes under different environments.

For simplicity we may assume that the four environmental conditions occur at equal probabilities (i.e., rainfall does not make the year cooler or vice versa). If we once again assume Hardy-Weinberg equilibrium and equal allele frequencies, the sexual population achieves a growth rate 22.5625 (1 − *s*) in every environmental setting, which also implies a geometric mean of 22.5625 (1 − *s*). The geometric mean for the asexuals is 16 for homozygote specialists (AABB, AAbb, aaBB, aabb), 18 for those who bet-hedge conservatively with respect to one trait only (AABb, aaBb, AaBB, Aabb), and 20.25 for the asexual genotype that conservatively hedges its bets with respect to both traits (AaBb). The sexual population can beat any asexual genotype if *s* < 0.1025, it can be beaten by the best bet-hedging asexual AaBb but not by others if 0.1025 ≤ *s* < 0.2022, it can be beaten by all bet-hedging asexuals (AABb, aaBb, AaBB, Aabb and AaBb) but beat the full homozygotes if 0.2022 ≤ *s* < 0.2909, and remains vulnerable to invasion by any asexual type if s exceeds 0.2909.

**Table 2:** Payoff of different genotypes under four different environmental conditions, when there are two traits impacting fitness.

| genotype | freq. | WH | WC | DH | DC |
| --- | --- | --- | --- | --- | --- |
| AABB | 1/16 | 64 | 16 | 16 | 4 |
| AABb | 1/8 | 32 | 32 | 8 | 8 |
| AAbb | 1/16 | 16 | 64 | 4 | 16 |
| AaBB | 1/8 | 32 | 8 | 32 | 8 |
| AaBb | 1/4 | 16 | 16 | 16 | 16 |
| Aabb | 1/8 | 8 | 32 | 8 | 32 |
| aaBB | 1/16 | 16 | 4 | 64 | 16 |
| aaBb | 1/8 | 8 | 8 | 32 | 32 |
| aabb | 1/16 | 4 | 16 | 16 | 64 |

We used specific numerical values in the example above, which raises the question how these generalise to other scenarios of allelic dominance, including dominance-recessive, heterosis and inbreeding depression. It has been shown that sexual population can reach all possible phenotypic states if and only if the hereditary system is either dominant-recessive or maternal or the combination of these (Garay and Garay, 1998). We show in appendix A that under the hereditary scheme where one allele is completely dominant over the other allele, the sexual heterozygote ceases to be a bet-hedging strategy since both its arithmetic mean and geometric mean fitness become equal to those of the asexual homozygote. Stronger dominance, on the other hand, improves the geometric mean fitness of the sexual population, making it potentially easier to outcompete asexuals.

## 3 Numerical simulations

In the previous section, we used the frequency distribution of different genotypes at Hardy-Weinberg equilibrium for calculating the arithmetic and geometric mean payoff of the sexual population. This is convenient, as it allows us to examine the situation as if the sexual population reached the same growth rate in every environmental setting (it makes sex achieve perfect bet-hedging in the sense that the geometric mean payoff equals the arithmetic mean payoff). However, in reality sex will fail to achieve this perfection, because the genetic environment encountered by a sexual population will be a function of past selection. There will then also be temporal variation in the distributions of genotypes, and sex is likely to fail to achieve perfect bet-hedging. The geometric mean fitness will then drop below the arithmetic mean fitness.

Since the pioneering work of Maynard Smith (Maynard Smith, 1971, 1976), Hamilton (Hamilton et al., 1981) and Bell (Bell, 1982), it has been known that the rate of temporal fluctuations can matter for the evolution of sex. In our setting above, the frequency of switches between wet and dry environments determines how far from equilibrium genotype frequencies will deviate over time. In the following we therefore use numerical simulations to show a more realistic picture of the competition dynamics between sexual and asexual populations.

### 3.1 Environmental fluctuations

Here we relax the assumption of Hardy-Weinberg equilibrium: it is only used as a starting state for sexual reproduction, and the following dynamics are computed according to a realised run of fluctuations of the environmental state. Assume that the wet and dry environments follow each other in a manner that can be captured by discrete-time Markov chains (i.e. the transition probability from one state to another does not depend on how long the environment has spent in the current state). The transition probabilities between states can be written in the matrix form

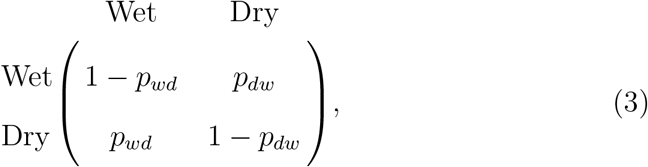

in which *p_wd_* denotes the probability that the environment changes from wet to dry in a year, and *p_dw_* is the probability that the environment changes from dry to wet in a year. The normalized dominant right eigenvector represents the stationary distribution of the environmental states (Caswell, 2001), and has the value (*p_wd_ /(*p*_*wd*_ + p_dw_); p_dw_ /(p_wd_ + p_dw_)*). The subdominant eigenvalue ρ = 1 − *p_wd_ − p_dw_* in turn corresponds to the correlation between the environmental states at times *t* and *t + 1* (Caswell, 2001). Therefore, consecutive environmental states are negatively autocorrelated if *ρ* < 0, positively autocorrelated if ρ > 0, and uncorrelated if ρ = 0. In the extreme case where *p_wd_ = p_dw_* = 1, we have ρ = −1 and wet and dry environments alternate, whereas in the other extreme case where *p_wd_ = p_dw_* = 0, we have ρ = 1 and the environment stays in the initial state forever.

### 3.2 Simulation results

To focus on the effect of environmental fluctuations, we exclude the effect of demographic stochasticity and drift by assuming that the population size is very large. We use the fixation probability of the invading type as a proxy for the relative advantages of different types. We do this by setting up a population consisting of an initial proportion 0.02 of the invading type, competing against one of the three possible alternative types. We assume that, for sexuals, the growth rate is proportional to 1 − *s* (the frequency of females), and the proportion of AA, Aa and aa young are derived by assuming that both male siring propensity and the female propensity to reproduce are proportional to that genotype’s payoffs (this covers at least two possible biological interpretations: survival probabilities are proportional to payoffs and thereafter mating is random, with each mating producing an equal number of offspring; or that the fecundity of females, as well as the siring success of males, is proportional to payoffs. As a caveat, note that the two cases can be mapped to each other directly only in unstructured populations. If the population has overlapping generations, selecting on survival and reproduction have to be treated separately from each other (Haccou and McNamara, 1998; Li et al., 2016)).

The invasion is tracked until one of three mutually exclusive events have happened: (a) the invading type has reached frequency 0.9999 or higher (we consider this a successful invasion, and fixation is reached), (b) the invading type’s frequency falls below 0.0001 (we assume that the invasion failed), or (c) neither (a) nor (b) have happened by generation 10^6^ (we consider this a coexistence scenario, but in practice event (c) never happened). The Octave codes for all numerical simulations are provided in Supplementary Information. The sexual population starts from the Hardy-Weinberg equilibrium state, with a proportion of 0.25 AA, 0.5 Aa and 0.25 aa types. The payoff of each genotype under different environments follows matrix (1), and fixation probabilities are estimated from 10^4^ independent realisations. Because the payoffs of the asexual AA and aa types are symmetric, and the wet and dry environments occur at equal frequencies, they have identical fixation probabilities when invading or being invaded by a sexual population. Therefore, without loss of generality, we use the asexual AA to represent the case of asexual homozygotes in Figure 1.

**Figure 1:**
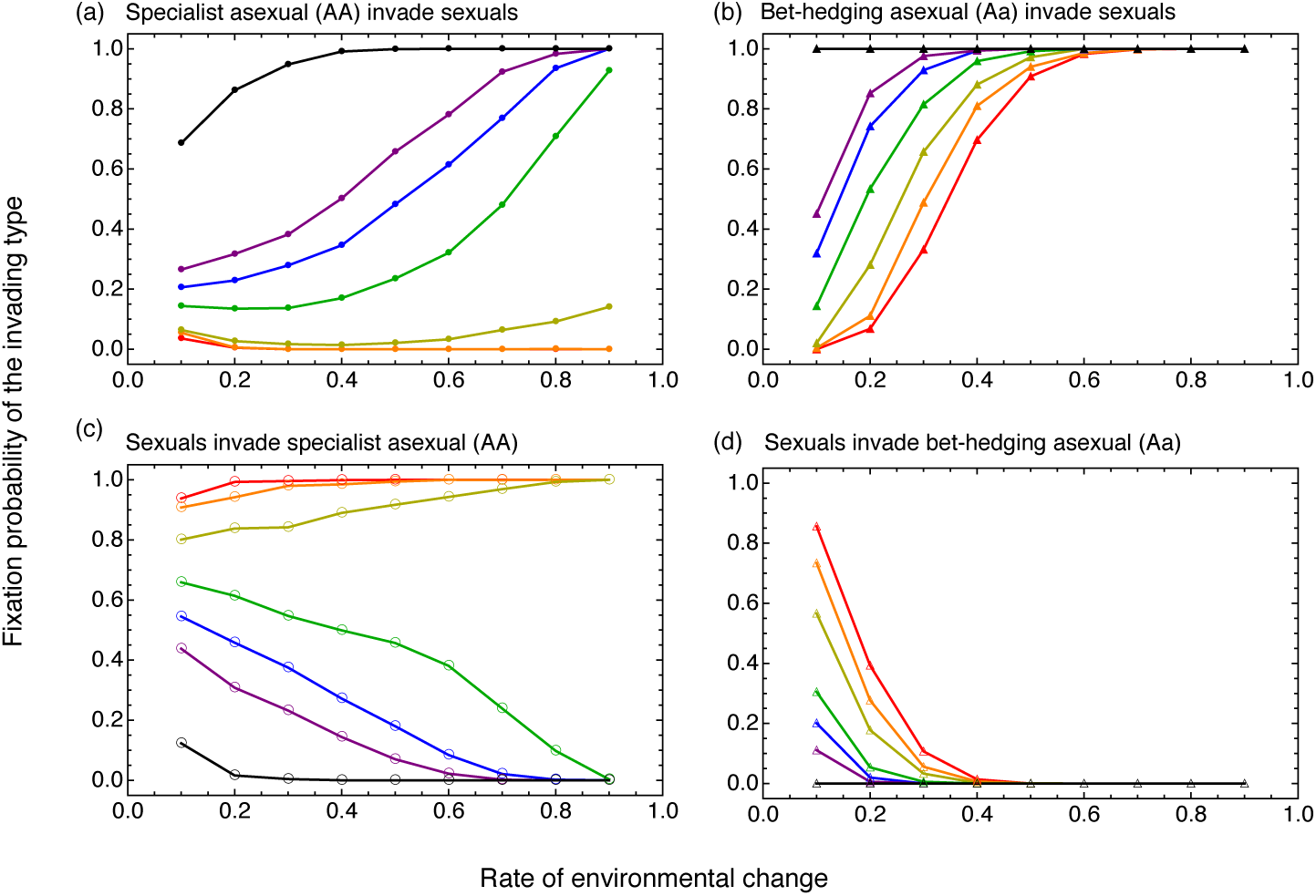
Fixation probability of the invading types under various rates of environmental change for populations following payoff matrix (1). The x-axis represents the rate of environmental change, assuming *p_wd_ = p_dw_*. Colours from red to purple to black represent sexual population of different sex ratios (0.01, 0.02, 0.04, 0.08, 0.12, 0.16, and 0.5). The larger the sex ratio, the higher the cost of sex. These figures are based on 10^4^ realisations per parameter value, and never required stopping the simulation at generation 10^6^ (i.e. either fixation is reached or the invader went extinct).

The figure confirms that sex has a difficult time invading asexual strategies if *s* = 0.5. If we elevate the chances for sexual reproduction to invade others by allowing *s* < 0.5, then cases where sex outcompetes specialist asex-uals (AA or aa) still typically do not predict that sex can also outcompete bet-hedging asexuals (comparing the left and right panels: curves are almost invariably higher on the right than on the left when considering an asexual invasion, and are always lower on the right than on the left when considering a sexual invasion). Whether fast or slow environmental fluctuations are best for sex is surprisingly complex. At very small *s*, sexuals are more likely to invade asexual homozygotes (and also resist their invasion attempts) if the environment changes fast. Other values of s predict the opposite. This complexity contrasts with early work on geometric mean fitness in the context of sex (Hamilton et al., 1981), predicting that a fast changing environment is beneficial to the maintenance of sex in general. But there are crucial differences between the payoff structures in his model and ours. (Note that Hamilton did not call Hamilton’s temporal fluctuation model bet-hedging).

The success of invasion is likely to depend on how long allelic diversity persists in the population. If the payoff of the heterozygote is low, and the environment changes relatively slowly, genetic diversity might become extinguished even before the asexual mutant is introduced. When the sexual population exists alone, it is possible that one allele, either a or A, is lost (examples: Figure 2a-b, mean time to extinction: Figure 2c-d). The better the heterozygote (Aa) payoff (Figure 2c), and the faster the environmental fluctuations (Figure 2d), the longer the coexistence time of both alleles. If one allele has already been lost, sex behaves genetically like an asexual homozygote (losing its bet-hedging benefit), but still paying the cost of sex. Note that a population that bet-hedges via asexuality (Aa) does not suffer from this risk, as both alleles are kept intact in this lineage in every generation. In this sense, conservative bet-hedging represented by asexuality may perform better than the diversified bet-hedging represented by sexual reproduction.

**Figure 2:**
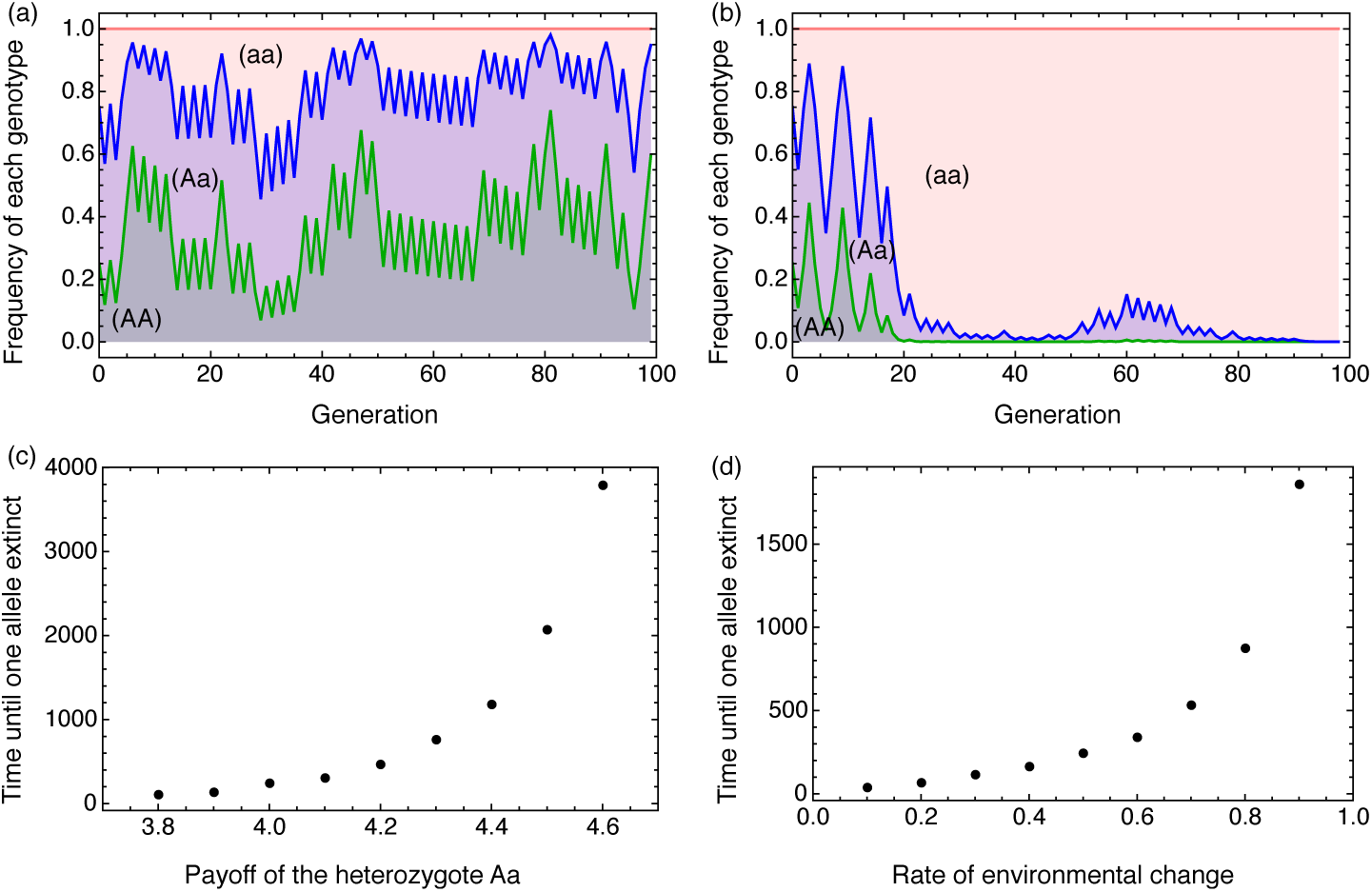
Examples of genetic diversity in a purely sexual population (no mutation to asexuality), where diversity is maintained (panel a) or lost (panel b) under environmental fluctuations that are tracked for 100 generations. The two trajectories are from simulations with identical parameter settings. In both cases, the rate of environmental change *p_wd_ = p_dw_* = 0.75, and the payoff of the heterozygote is set to 3.8 under both environmental conditions. The vertical height of regions of various colours represent the frequencies of different genotypes. (c) The mean time to the disappearance of one allele as a function of varying heterozygote payoffs when *p_wd_ = p_dw_* = 0.5, and (d) the mean time to the disappearance of one allele as a function of the rate of environmental change when *p_wd_ = p_dw_* and heterozygote payoff is 4.0 under both environmental conditions. In all simulations, the payoffs of the homozygotes follow payoff matrix (1). In panels (c) and (d), one allele is considered to have gone extinct if the frequencies of both the corresponding homozygote and the heterozygote are smaller than 10^−4^.

**Figure 3:**
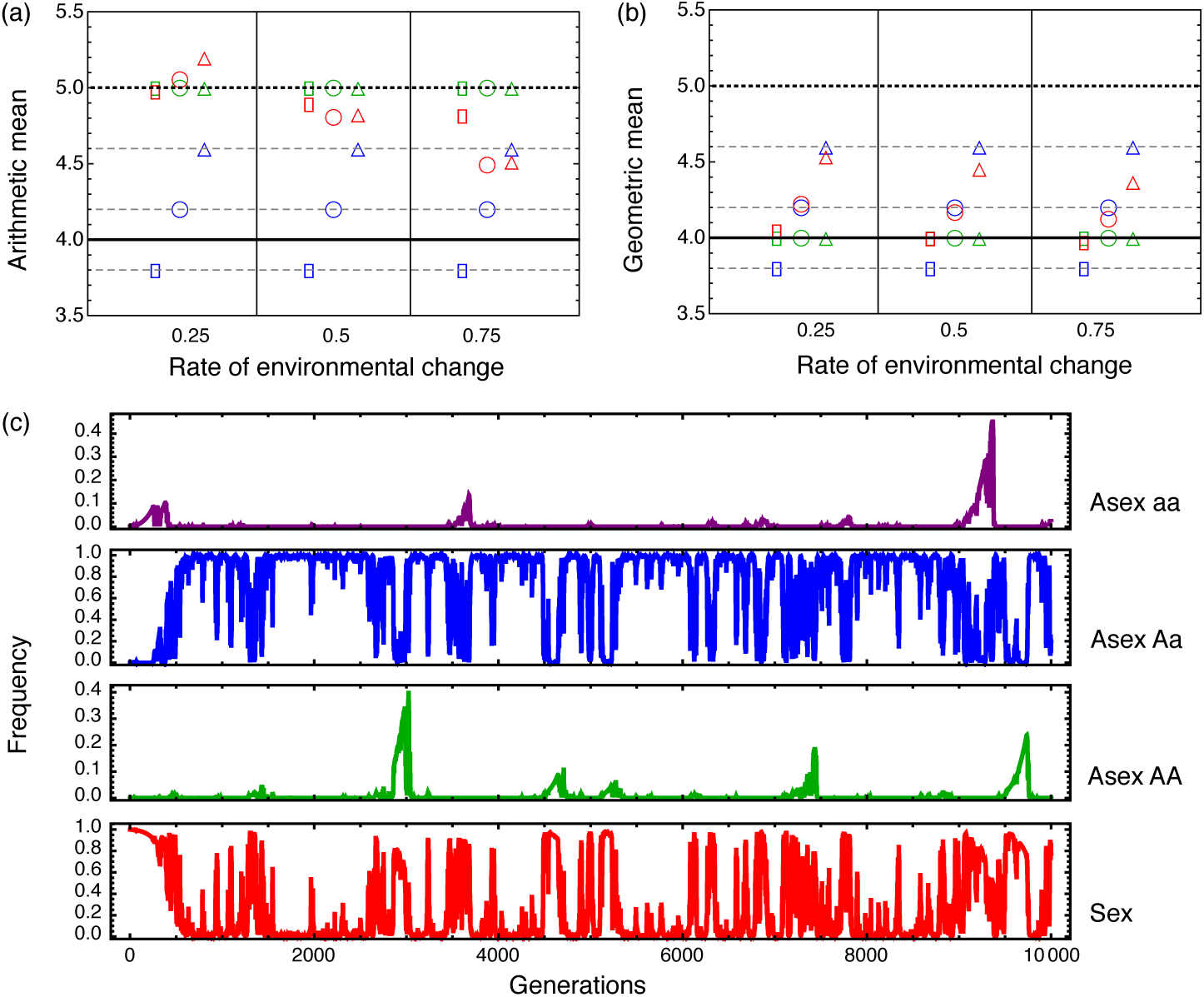
(a) Arithmetic mean payoffs and (b) geometric mean payoffs of the asexual homozygote (green), asexual heterozygote (blue) and the sexual population (red), computed over 500 generations when the payoffs of the asexual homozygotes follow matrix (1) and the sex ratio of the sexual population is set to *s* = 0.01. Symbols of different shapes represent different payoffs of the heterozygote: square, circle and triangle stand for 3.8, 4.2, and 4.6 respectively. The black dotted line is the expected arithmetic mean payoff of the asexual homozygotes, the black solid line is the expected geometric mean payoff of the asexual homozygotes, and the grey dashed lines are the expected arithmetic and geometric mean payoff of the asexual heterozygote. (c) Frequency dynamics of the sexual population and each asexual genotype under a changing environment over 10000 generations. In each panel, the x-axis is time (the elapsed number of generations), and the y-axis is the frequency of each type. All four panels are from the same instance of simulation. The heterozygote payoff is set to 4.2, and the rate of environmental change is *p_wd_ = p_dw_* = 0.5. The simulation starts with a pure sexual population with 0.25 AA, 0.5 Aa and 0.25 aa genotypes, but each individual may mutate to being asexual if previously sexual, or sexual if previously asexual, at rate 0.0001 per generation.

A key finding is therefore that sex cannot easily outcompete asexual forms based on bet-hedging benefits alone (Figure 3). Sex as bet-hedging requires conditions under which the red symbols are below the dotted line in Figure 3a, and above the solid line in Figure 3b. Only four out of the nine cases satisfy the requirements (heterozygote payoff 4.2 or 4.6 in combinations with rate of environmental change 0.5 or 0.75). However, it is possible to construct cases where sex wins in terms of arithmetic mean fitness but loses in terms of geometric mean to the conservative asexual bet-hedger (Figure 3c, where the heterozygote payoff is set to 4.2, and the rate of environmental change is set to 0.5).

## 4 Discussion

There are interesting parallels between sex and bet-hedging theory. Intuitively, the costs of sex reduce the fitness of sexual lineages in every generation that undergoes a sexual life cycle (hence the arithmetic fitness is reduced), but by diversifying the genotypes of offspring, sex can reduce the variance in success: in any given year some offspring will survive, while an asexual specialist proverbially puts “all its eggs in one basket” – leading to very low success if the year features a mismatch between offspring genotype and the state of the environment. However, for this to favour sex over asex, the geometric mean fitness of the former should be elevated above the latter. Although variance reductions have a beneficial effect on geometric mean fitness, arithmetic mean fitness (which is low for sexual types) simultaneously sets an upper limit for it, and hence it is not easy for sex to reach such high bet-hedging benefits that its geometric mean fitness is the best of all competing strategies. In other words, the fact that sexual reproduction shows features of bet-hedging is not the same statement as the claim that bet-hedging provides strong enough benefits for the evolution and maintenance of sex. This is especially true since sex may have to compete against another type of bet-hedger: that of asexual heterozygotes, which avoid paying the cost of sex but may also achieve bet-hedging if their genotype performs reasonably well under all considered environmental conditions. This highlights that (a) it is important to specify that a strategy is performing bet-hedging relative to another strategy, and be explicit about the identity of the relevant competitor, and (b) that it would be premature to consider bet-hedging as a major driving force behind the maintenance of sex, at least under the simplifying assumptions of the current model.

Fast and unpredictable changes of the environment have been found to favour bet-hedging (Haccou and Iwasa, 1995) and facilitate the maintenance of sexual reproduction (Maynard Smith, 1971, 1976; Treisman, 1976; Hamilton et al., 1981; Bell, 1982; Waxman and Peck, 1999; Barbuti et al., 2012), but these authors did not use bet-hedging terminology. Our model shares a similar genetic structure to Hamilton et al. (1981), but the payoff structures are different. In our model, the two asexual homozygotes are specialists that adapt to different environmental conditions, and the heterozygote has intermediate payoff under both environmental conditions (this makes it a conservative bet-hedger). In Hamilton’s model, the homozygotes receive identical payoffs (that depend on environmental conditions), whereas the payoff of the heterozygote is the reciprocal of this payoff. The heterozygote and ho-mozygotes in the model of Hamilton et al. (1981) thus do not correspond to a bet-hedger and two specialists, and therefore, although the model shows that sex is beneficial under a fast changing environment, it did not aim to capture the evolutionary dynamics under the bet-hedging context.

Compared to classic bet-hedging scenarios where the bet-hedger always has the same payoff under the same environment (Starrfelt and Kokko, 2012), sexual reproduction as bet-hedging brings in additional features. In the sexual population, the arithmetic mean payoff in each generation is determined not only by the environment, but also the frequency distribution of all genotypes, the sex ratio, and possibly other costs or benefits from sexual reproduction. In addition, if mutations between sexual and asexual populations are allowed, more than one type of bet-hedging strategy can (at least temporarily) coexist, and it is insightful to remember that there can be asexual heterozygotes that bet-hedge conservatively, as opposed to the diversified bet-hedging of the sexual population.

Both theoretical and experimental work on the evolution of sex show complications that highlight the simplicity of any two-environment model (indeed, in our model too, increasing the dimensionality of the system helps maintain sex). We have followed a tradition in bet-hedging theory where 2 (or 4) types of environment can be adapted to with one (or two) traits. Modern research on genetic variation reveals that there is surprisingly much polygenic variation present in populations (Charlesworth, 2015), and fitness landscapes are often complex. Recent research on sex has revealed the potential importance of processes such as clonal interference (McDonald et al., 2016; Sharp and Otto, 2016), which tends to erode the success of asexual lineages over time because they are slow to acquire multiple novel mutations that aid adaptation. Sex improves the rate with which innovations end up in the same organism, while asexual lineages tend to fail in having access to the most “up to date” genetic background, especially if the environment keeps changing. The detrimental interference between competing clones that have acquired one or another beneficial allele (at different loci) eventually makes asexuality an inferior competitor in the adaptive race. While this is a very different situation from what bet-hedging theory traditionally has considered, there is scope to fill this gap: the gist of the argument is that the asexual lineages experience diminishing geometric fitness once timescales become long enough that novel beneficial mutations begin playing a role. Sex and the diversity it creates can help diversify the genetic backgrounds where new mutations can be selected for.

Among the classic literatures, the payoff structure in Treisman (1976) is the closest to ours, and it also captures some of the above ideas about the environment changing to something never experienced before. In Treisman (1976), different alleles interact additively and give the diploid individual a phenotype (in his words, a “genotypical score”) that impacts female fertility but not male siring success. Alleles have effects of −0.5 or 0.5, so that homozygotes have phenotypes –1 or 1, and the heterozygote has an intermediate phenotype of 0. Females (both sexual and asexual) can only breed if their phenotype matches, within tolerable range, the environmental conditions (such as temperature). If the environment keeps changing (e.g., increasing temperatures), asexual genotypes cannot keep pace with sexuals that produce diversified offspring through recombination; asexual extinction can then follow. Treisman (1976), like the authors mentioned above, did not use the terminology of bet-hedging, and hence did not analyse the arithmetic and geometric mean fitness of each genotype.

Given that there is both old and new work on sex that could gain conceptual clarity if researchers routinely reported how the winning strategy (sexual or asexual) performed in terms of arithmetic and geometric mean fitness, we welcome more work in the areas linking sex and bet-hedging. Bet-hedging theory has brought about increased understanding of other evolutionary questions from dispersal evolution (Armsworth and Roughgarden, 2005) and dormancy timing (Ellner, 1985; Evans and Dennehy, 2005; Furness et al., 2015) to antibiotic resistance (Arnoldini et al., 2014), microbial population dynamics (de Jong et al., 2011) and phenotypic switching (Carja et al., 2014). It would appear timely to add sexual reproduction to this list. Even if sex in simplistic settings (like ours) does not reach the status of a strategy with the highest geometric mean fitness, a bet-hedging perspective can shed light on the precise reasons why it failed. An interesting question would be to use this type of analysis to examine cases where sex, e.g. in situations involving clonal interference and *de novo* mutations, succeeds to maintain itself against asexual competitors.

## Acknowledgement

We thank the two anonymous reviewers for their comments and suggestions. X.L. and H.K. are grateful to the Swiss National Science Foundation. J.L. was funded by a University of New South Wales Vice-Chancellor’s Post-doctoral Research Fellowship. All authors thank the organisers of the 17th International Symposium on Dynamic Games and Applications.

### A Sex as bet-hedging when one allele dominates the other

Assume that the *A* allele fully dominates the *a* allele. The fitness values of each genotype under different environments are show in matrix (4).

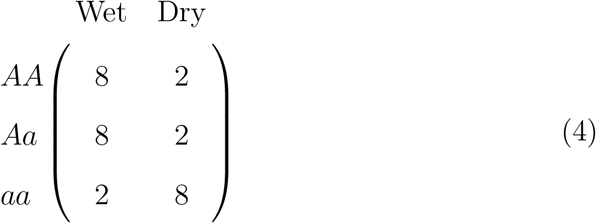

In this case, the payoff of each asexual type and the sexual population is shown in table 3.

**Table 3:** The payoff structure under wet and dry years when the A allele fully dominates the a allele: the arithmetic mean (AMean) and the geometric mean (GMean) of the payoffs of asexual lineages, as well as of a sexual population assumed to be at the Hardy-Weinberg equilibrium.

|  | Wet | Dry | AMean | GMean |
| --- | --- | --- | --- | --- |
| asex-$AA$ | 8 | 2 | 5 | 4 |
| asex-$Aa$ | 8 | 2 | 5 | 4 |
| asex-$aa$ | 2 | 8 | 5 | 4 |
| sex-population | $6.50(1-s)$ | $3.50(1-s)$ | $5(1-s)$ | $4.77(1-s)$ |

The first observation is that the asexual heterozygote is no longer a bet-hedging strategy, since its payoffs under different environmental conditions become identical to the homozygote *AA*, and thus its geometric and arithmetic payoffs no longer fit the requirements of bet-hedging. Under Hardy-Weinberg equilibrium, the sexual population would have higher geometric mean payoff and lower arithmetic mean payoff than each asexual type when 0 < *s* < 0.162. This range is larger than that under the case of intermediate inheritance, where the sexual population beats any asexual homozygote if 0 < *s* < 0.158, and beats the asexual heterozygote if 0 < *s* < 0.053.

Similar results hold when populations hedge their bets on multiple traits. Using the case in matrix (2) as an example, if the *A* allele fully dominates the a allele, and the *B* allele fully dominates the b allele, the payoff matrices for rainfall and temperature adaptation has the following structure:

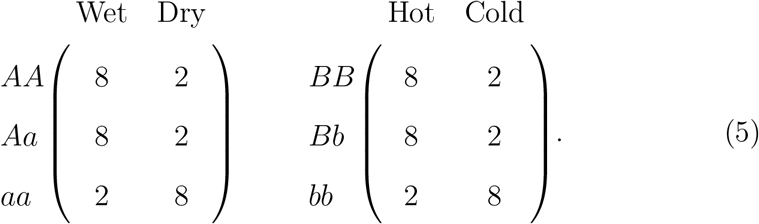

Again, we assume that different traits interact multiplicatively to determine the final fitness, and the sexual population is under Hardy-Weinberg equilibrium. Table 4 gives the complete list of payoffs of different genotypes under different environments.

**Table 4:**
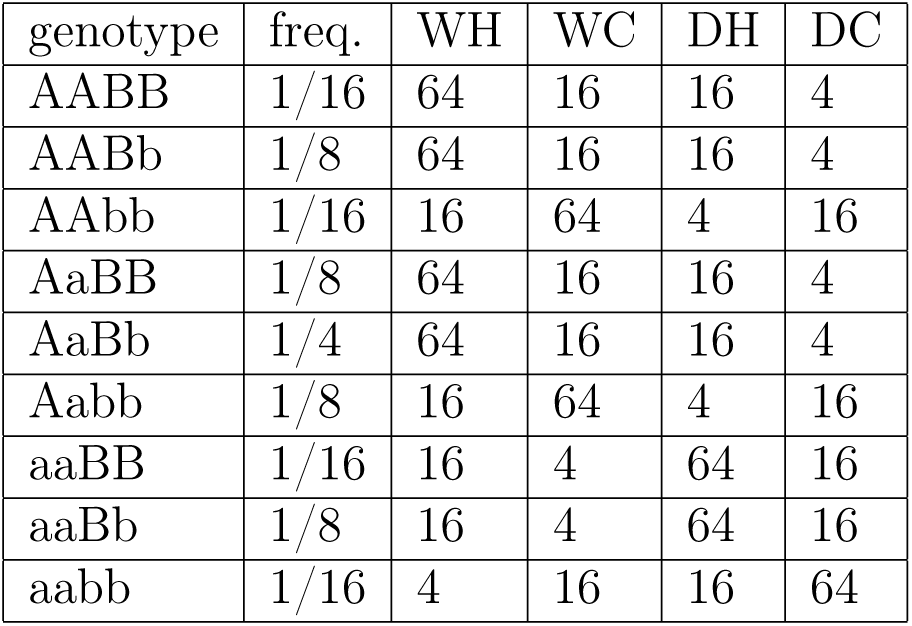
Payoff of different genotypes under four different environmental conditions under the dominance hereditary system, when two traits determine the fitness together.

| genotype | freq. | WH | WC | DH | DC |
| --- | --- | --- | --- | --- | --- |
| AABB | 1/16 | 64 | 16 | 16 | 4 |
| AABb | 1/8 | 64 | 16 | 16 | 4 |
| AAbb | 1/16 | 16 | 64 | 4 | 16 |
| AaBB | 1/8 | 64 | 16 | 16 | 4 |
| AaBb | 1/4 | 64 | 16 | 16 | 4 |
| Aabb | 1/8 | 16 | 64 | 4 | 16 |
| aaBB | 1/16 | 16 | 4 | 64 | 16 |
| aaBb | 1/8 | 16 | 4 | 64 | 16 |
| aabb | 1/16 | 4 | 16 | 16 | 64 |

In this case the sexual population has a fitness of 42.25 (1 – *s*) under the WH environment, 12.25 (1 − *s*) under the DC environment, and 22.75 (1 − *s*) under both WC and DH environments. Therefore, if four different environments occur at equal frequencies, the arithmetic mean payoff of the sexual population is 25 (1 − *s*), and the geometric mean fitness is 22.75 (1 − *s*). The geometric mean for the asexuals is 16 for all asexual types. In this way, the sexual population beats any asexual population if 0 < *s* < 0.297. This range is also larger than the condition (0 < *s* < 0.102) for beating any asexual genotype under the intermediate heredity.

